# Purkinje cell number-correlated cerebrocerebellar circuit anomaly in the valproate model of autism

**DOI:** 10.1101/434217

**Authors:** Tamás Spisák, Viktor Román, Edit Papp, Rita Kedves, Katalin Sághy, Cecília Katalin Csölle, Anita Varga, Dávid Gajári, Gabriella Éva Nyitrai, Zsófia Spisák, Zsigmond Tamás Kincses, György Lévay, Balázs Lendvai, András Czurkó

## Abstract

While cerebellar alterations may play a crucial role in the development of core autism spectrum disorder (ASD) symptoms, their pathophysiology on the function of cerebrocerebellar circuit loops is largely unknown. We combined multimodal MRI (9.4 T) brain assessment of the prenatal rat valproate (VPA) model and correlated immunohistological analysis of the cerebellar Purkinje cell number to address this question. We hypothesized that a suitable functional MRI (fMRI) paradigm might show some altered activity related to disrupted cerebrocerebellar information processing. Two doses of maternal VPA (400 and 600 mg/kg, s.c.) were used, and while the higher VPA dose induced a global decrease in whole brain volume, the lower dose induced a focal gray matter density decrease in the cerebellum and brainstem. Increased cortical BOLD responses to whisker stimulation were detected in both VPA groups, but it was more pronounced and extended to cerebellar regions in the 400 mg/kg VPA group. Immunohistological analysis revealed a decreased number of Purkinje cells in both VPA groups. In a detailed analysis, we revealed that the Purkinje cell number interacts with the cerebral BOLD response distinctively in the two VPA groups that highlights atypical function of the cerebrocerebellar circuit loops with potential translational value as an ASD biomarker.

## Introduction

Autism spectrum disorder (ASD) is a lifelong neurodevelopmental disorder ^1^, currently diagnosed using behavioral tests that can be subjective; therefore, objective noninvasive imaging biomarkers of autism are being actively researched ^2–5^.

Cerebrocerebellar circuit dysfunction may play a crucial role in the etiology of ASD, as cerebellar lesions or structural and functional differences in various cerebellar subregions consistently have been linked to ASD ^6–11^. MRI findings of the cerebellum in ASD, however, remain less clear ^12^: although the size of the vermis appears to be slightly smaller and structural differences in cerebellar lobule VII (right crus I/II) are associated with core ASD symptoms and may have potential functional impact ^6, 7^. Similarly, the great majority of a substantial number of postmortem anatomical studies of ASD show decreased Purkinje cell density, but with focal regional alterations ^6, 13^.

Neuroanatomical studies underline the extensive and region-specific connectivity between the cerebellum and the cerebral cortex with two-stage feed-forward and feed-back loops, e.g., lobules VI/VII (crus I/II) of the cerebellum are linked with the association areas of the cerebral cortex concerned with higher order behavior, such as the prefrontal cortex, posterior parietal cortex or cingulate gyrus ^14, 15^. The anterior part of the cerebellum has a sensory domain, in addition to its traditionally known role in motor functions, and the posterior part of it contributes to some aspects of affective and cognitive processing ^16–18^. Indeed, cerebellar posterior lobe lesions define cerebellar cognitive affective syndrome (CCAS), and *in utero* or early postnatal lesions induce behavioral and social deficits that overlap with ASD symptoms ^15, 19^.

Despite these seminal findings highlighting the potential functional impact of disrupted cerebrocerebellar loops in ASD ^8, 20^, fundamental questions about the pathophysiology of these cerebrocerebellar loops have remained unanswered. Equally, their translational utility is unknown, limiting the progress in pursuing objective neuroimaging biomarkers in ASD.

We addressed these questions by combining multimodal neuroimaging and correlated immunohistological analysis of the cerebellar Purkinje cell number in a prenatal rat valproate (VPA) model. We hypothesized that our functional MRI (fMRI) paradigm might show some altered activity related to disrupted cerebrocerebellar information processing. We used a somatosensory functional fMRI paradigm suitable for small animal imaging. Our paradigm, by involving nested loops of the whisker system at the cerebellar, midbrain and thalamocortical level ^21^, allocates reasonable translational power to sensory-attentional paradigms in human studies. To ensure that our findings are unspecific to any developmental phase, we performed an MRI assessment of the same cohort twice, with a one-month difference (measurement 1 and 2). Furthermore, due to the uncertainty regarding the optimal maternal VPA dose, in addition to a control group with maternal vehicle treatment, we investigated two VPA groups, with maternal doses of 400 mg/kg and 600 mg/kg (hitherto referred to as the VPA400 and VPA600 groups).

## Results

### Valproate-induced reduction of cerebellar gray matter and whole brain volume

Because both human ASD and its VPA model are known to present specific morphological features, our first assessment was to perform volumetry and voxel-based morphometry (VBM) analysis on the structural MRI scans. Anatomical scans at measurement 1 revealed that prenatal exposure to VPA resulted in 3.2% smaller whole brain volume in the VPA600 group (p=0.006, effect size: −81.1 mm^3^) compared to controls. Smaller brain size was also observed in gray matter (−3.2%, p=0.02, effect size: −52.7 mm^3^) and white matter (−2.4%, p=0.09, −13.2 mm^3^) volume; however, the latter was not significant. A milder difference in the whole brain volume (−2%, p=0.06, effect size: −51.9 mm^3^) was observed in the VPA400 group, and this was not detectable either in the gray or in the white matter volume. Detailed summary statistics of the volumetry data are listed in Supplementary Table 2.

VBM analysis of the structural MRI scans was used to localize the whole brain averaged gray matter volumetric differences. In the VPA600 group, the VBM analysis could not detect any well-localized source of the smaller global gray matter volume; there were no normalized voxels showing significantly lower gray matter density (signal intensity modulated by Jacobian determinants) compared to the vehicle group. On the other hand, significantly lower focal gray matter density was observed in the VPA400 group in various brainstem nuclei and the bilateral cerebellar crus I/II and paraflocculus compared to controls (Fig. 1). Comparing the VPA400 and VPA600 groups revealed a highly similar pattern of localized focal hypoplasia in the VPA400 group, suggesting that this cerebellar and brainstem gray matter difference is specific to the 400 mg/kg prenatal dose. No positive differences in gray matter density were observed in either of the VPA groups compared to the VEH group.

**Figure 1.**
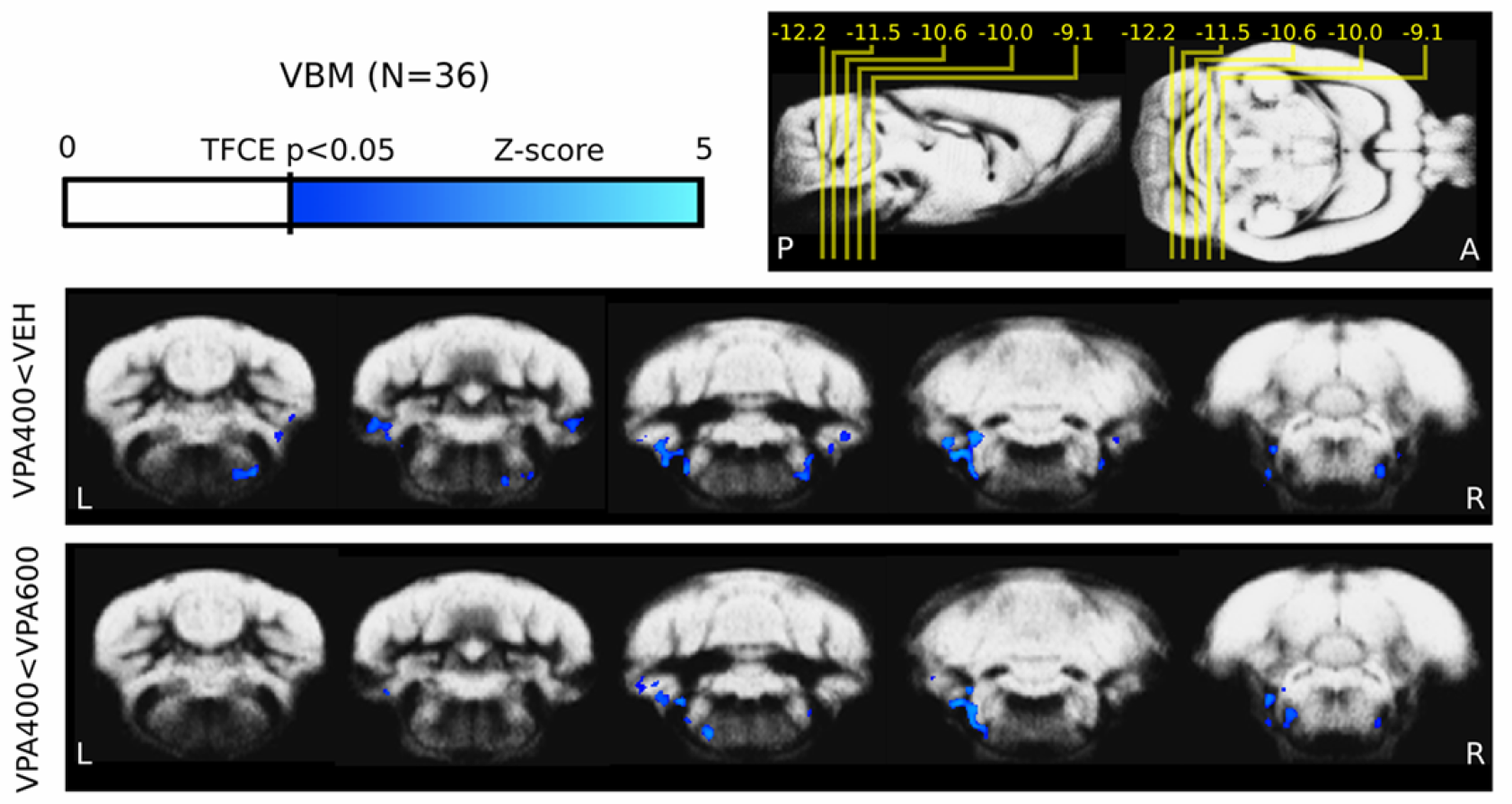
Decreased cerebellar and brainstem gray matter density revealed by voxel-based morphometry analysis. Evidence of cerebellar (crus I/II and paraflocculus) and brainstem (lemniscus) gray matter hypoplasia was observed only in the VPA400 group compared to both the VEH (top) and the VPA600 (bottom) groups (measurement 1). Compared to the VEH group, no decreases in VPA600 group and no increases in either of the VPA groups were observed. These results, together with the tissue volumetry analysis (Fig. 3), suggest that the 400 mg/kg VPA dose might be in an appropriate range for triggering autistic-like morphological features, as opposed to the 600 mg/kg dose, which in turn exhibits cortical hypoplasia without cerebellar alteration. On the top right of the figure, the position of slices (and their distance from bregma in mm) are shown on sagittal and horizontal planes. The colorbar represents Z-score (Gaussianized T-score) values of the parametric images (TFCE enhanced FDR-corrected significance threshold of p=0.05). The slices highlight all clusters of significant activations (meaning that there are no significant changes in the cerebrum). Parametric images are overlaid on the study specific gray-matter probability map. Images are shown in neurological convention (left-is-left; A: anterior, P: posterior, L: left, R: right)

### BOLD response to whisker stimulation

We assessed the possible ASD-related functional changes in the VPA rat model in the sensory domain by somatosensory stimulation-based functional MRI. Whisker stimulation evoked a widespread, significant BOLD response in all three animal groups (Supplementary Tables 3-5, Fig. 2A). As expected, the activation peaked in the barrel field of the primary somatosensory (S1) cortex (contralateral to the stimulation side) and in a blob covering the primary auditory (Au1) and the secondary somatosensory (S2) cortices. The group-mean maximal signal change in S1 was 0.36%, 0.68% and 0.34% in the VEH, VPA400 and VPA600 groups, respectively. The ipsilateral somatosensory activation was weaker but also significant in all three groups. Additionally, bilateral striatal, frontal, orbitofrontal, insular, entorhinal, cerebellar, and amygdalar as well as brainstem (trigeminal) activations were found in all three groups. Local maxima and effect sizes are reported in Supplementary Tables 3-5. In general, group-level activation was the most extended in the VPA400 group (455.8 mm^3^), followed by the VPA600 (225.2 mm^3^) and the vehicle (190 mm^3^) groups.

**Figure 2.**
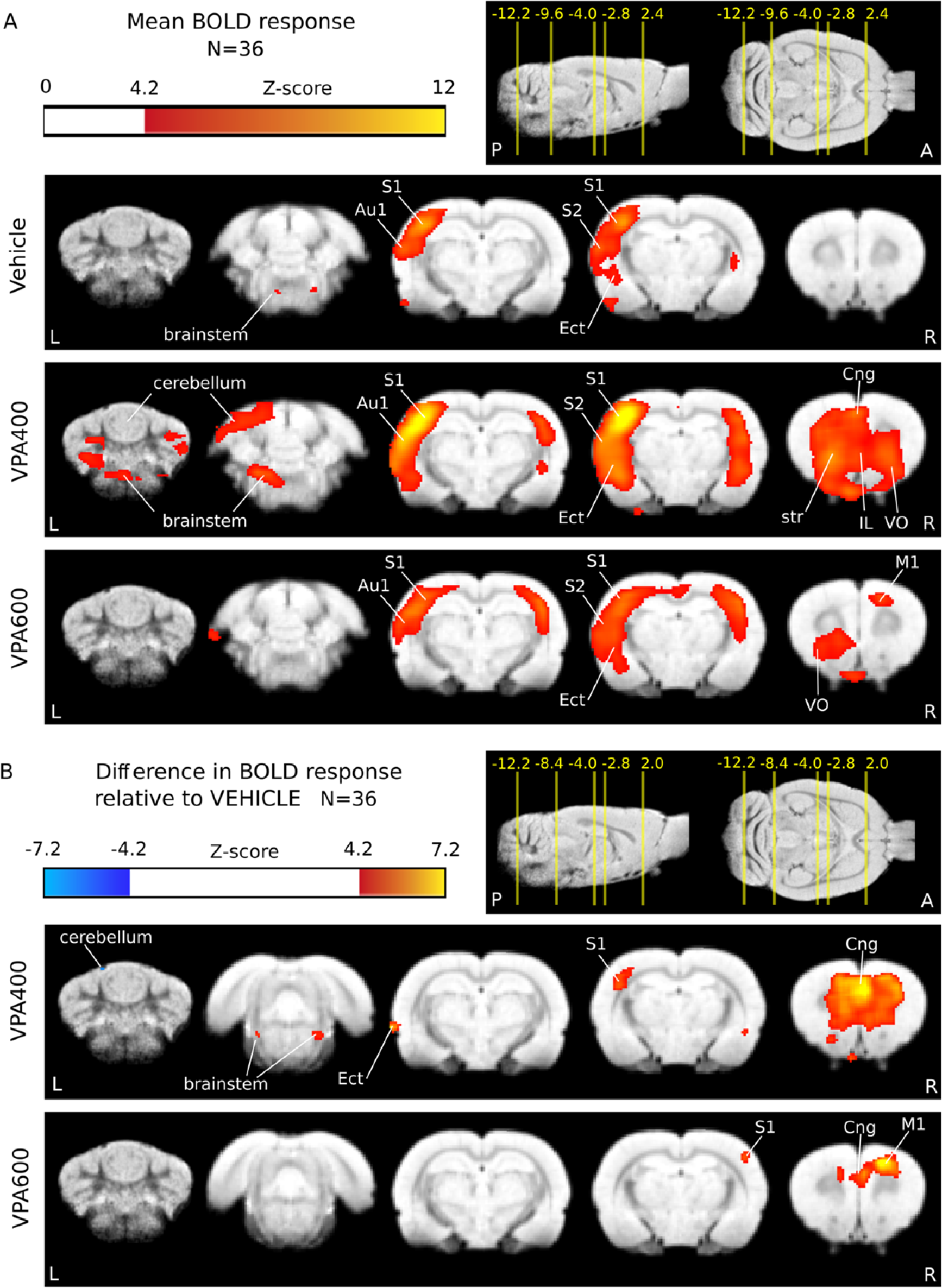
BOLD hyperactivation in response to somatosensory stimulation after prenatal valproate exposure. Whisker stimulation triggered activation mainly in the contralateral primary and secondary somatosensory cortices (S1 and S2, respectively), auditory cortices (Au1) and the trigeminal nuclei of the brainstem. (B) BOLD response differences in the VPA400 and VPA600 groups compared to the VEH control group showed a robust activation increase in S1, anterior cingulate cortex (Cng), frontal areas (IL: infralimbic cortex, VO: ventral orbital cortex) and brainstem lemniscus in the VPA400 group. A spatially restricted decrease was also observed on the cerebellar surface in this group. A similar but less prominent frontal and ipsilateral somatosensory hyperactivation was observed in the VPA600 group and involved the primary motor cortex (M1). Activations are displayed on coronal slices, overlaid on an in-house standard proton-density template. On the top right of each panel, the position of the slices are displayed on sagittal and horizontal planes as a distance from bregma in mm. Colorbars represent Z-score values. Significance thresholding was corrected for multiple comparisons (p=0.05). Abbreviations: A: anterior, P: posterior, L: left, R: right.

### Functional hyperactivation as a consequence of prenatal valproate exposure

Between-group fMRI analysis contrasts revealed that several of the striking differences of the groupwise activation patterns were statistically significant (Fig. 2B, Table 1). A remarkable and robust effect of prenatal VPA exposure was the widespread increase in the BOLD response to whisker stimulation (Fig. 2B, Table 1). In both the VPA400 and VPA600 groups, pronounced hyperactivation was found in the primary somatosensory, ventral orbital, anterior cingular and frontal association cortices. No decreased activation was detected in either of the VPA groups.

**Table 1.**
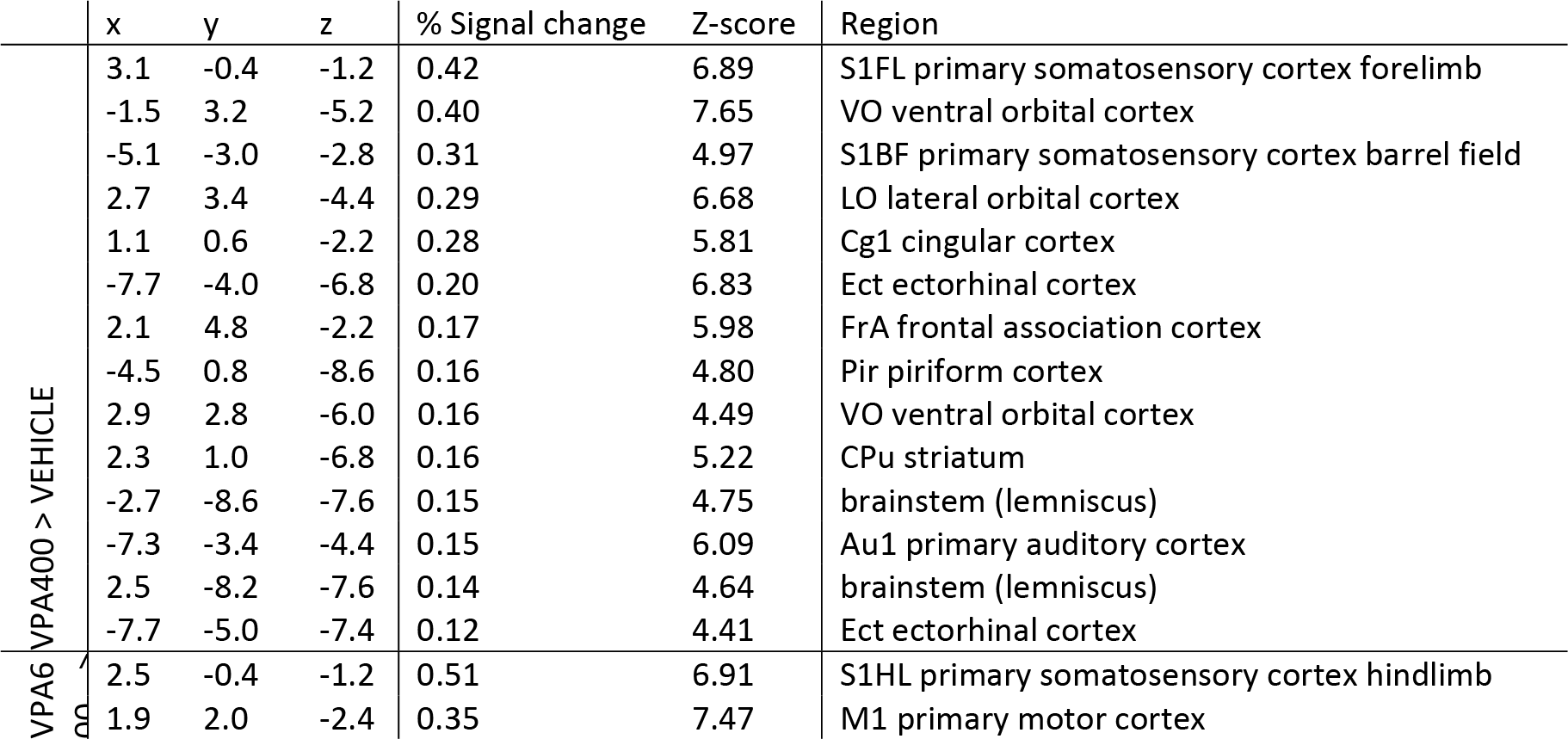
Local maxima of the change of the fMRI BOLD response in the VPA400 and VPA600 groups compared to the VEH control group. Only local maxima above the FWER P>0.05 threshold (|Z|>4.29) are shown. The minimum peak distance is set to 3 mm. Peaks are listed by effect size (% BOLD signal change) in decreasing order. Coordinates are given relative to bregma, according to the atlas of Paxinos and Watson (x: medial-lateral, y: anterior-posterior, z: dorsal-ventral).

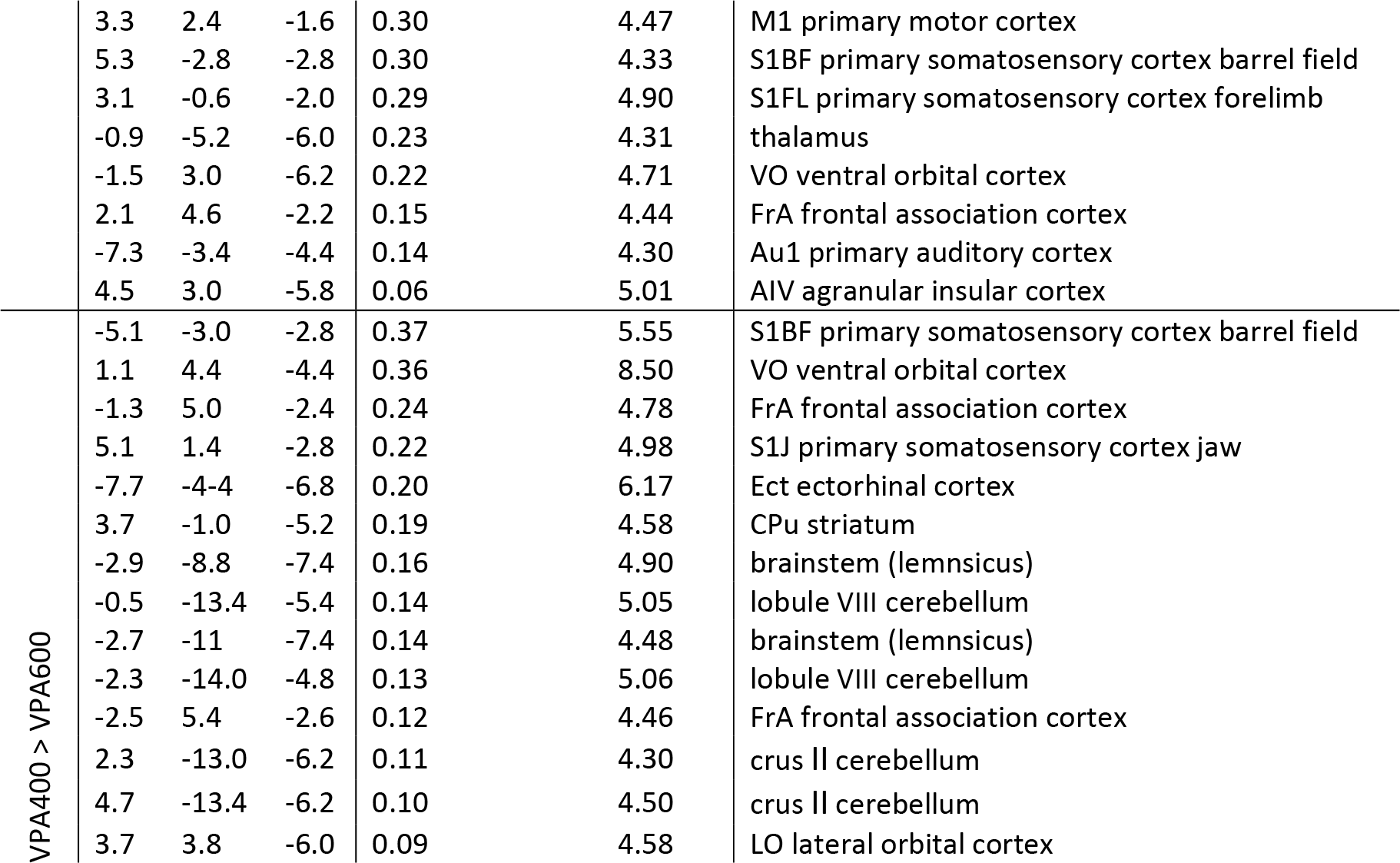

Contrasting the VPA400 and VPA600 groups revealed stronger hyperactivation in the VPA400 group, especially in regions of the primary somatosensory (contralateral to stimulation), ventral orbital, and frontal association cortices (Table 1) but also in the cerebellar lobule VIII and crus I/II. (Table 1).

### Repeated MRI assessment of the functional and structural alterations

Measurement 2 (performed one month later) reproduced the results of measurement 1, indicating that most of our MRI findings persevered (Fig. 3C-F).

**Figure 3.**
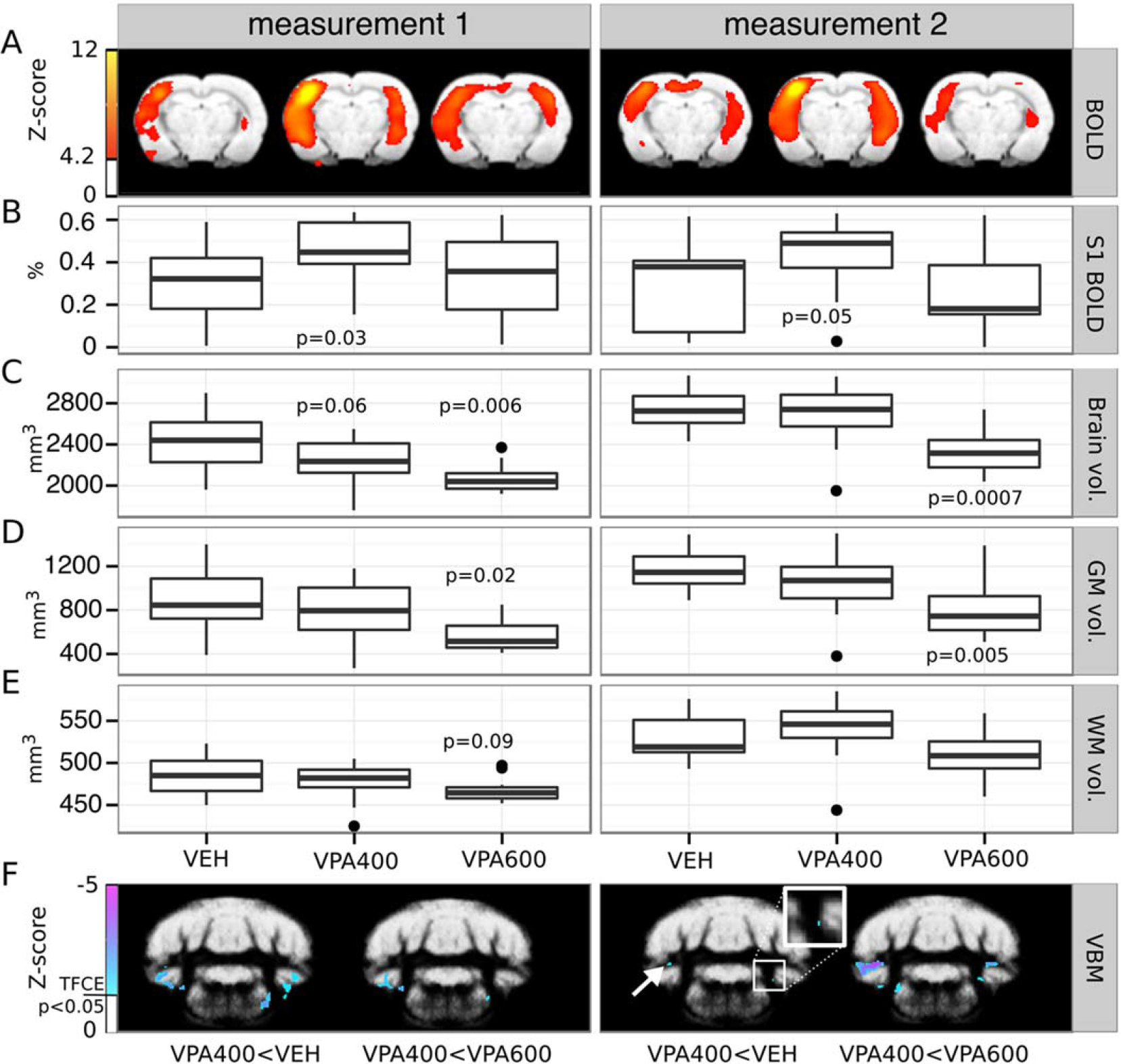
Persevered prenatal valproate-induced alterations demonstrated by a repeated MRI assessment of brain structure and function. (A) fMRI statistical parametric maps (at a coronal slice 2.8 mm from bregma in the posterior direction) of group-mean BOLD responses to whisker stimulation in the three experimental groups measured with one month difference. (B) Boxplots represent the average BOLD response in an unbiased somatosensory ROI (pooled-group mean activation Z-score > 9). Note the reproducible hyperactivation in the contralateral somatosensory cortex in the VPA400 group, in both measurements. (C, D, E) Boxplots depict the volume (mm^3^) of the whole brain, gray matter and white matter, respectively. The VPA600 group exhibits a significantly decreased whole brain volume and gray matter volume in both measurements, despite the overall increase in brain volume. (F) Bilateral cerebellar and brainstem gray matter density decreases found by VBM in the VPA400 group compared to both the VEH group (left image in both panels) and the VPA600 group (right image in both panels), consistently in both MRI measurements. Inset zooms into small details of the image, revealing a spatially limited cluster of difference. Colorbars represent Z-score values. Statistical images are thresholded at a corrected threshold of FWER P<0.05. Images are displayed in neurological convention (left-is-left). Boxes in the boxplots represent the 75th, 50th (median), and 25th percentile, whereas whiskers represent either the extrema or twice the interquartile range. Observations falling outside the 2*IQR range are represented by points. P-values lower than 0.1 (contrasting VPA groups to the VEH) are shown at the boxplots.

Despite the overall increase in the brain volume of all groups (+3.2%, from 2564.8 mm^3^ to 2646.4 mm^3^ in the VEH group, see Supplementary Table 1), a 3.6% difference between the VPA600 and the VEH group still persisted (p=0.0007, effect size: −97.9 mm^3^). This difference was also present in gray matter (−3.3%, p=0.005, effect size: −54.5 mm^3^).

The VBM analysis of measurement 2 demonstrated decreased gray matter density in cerebellar and brainstem locations of the VPA400 group with a remarkable overlap with measurement 1 (Fig. 3F). The decrease was also observed in contrast to the VPA600 group, while there was no change in gray matter density in any other group contrast.

In measurement 2, the group-level spatial pattern of whisker stimulation-induced BOLD activations were similar to those of measurement 1 (Supplementary Fig. 1.), although with smaller magnitudes due to the higher isoflurane doses. This and the smaller sample size due to exclusions (e.g., in-scanner motion; N=33) resulted in a decreased statistical power. Nevertheless, we were still able to detect functional hyperactivation in the VPA400 group compared to the vehicle group. The hyperactivation was significant in a cluster in the primary somatosensory cortex (contralateral to stimulation, cluster-level correction for multiple comparisons, P_clust_=0.045, clusterwise mean increase in effect size: 0.28%, see Supplementary Fig. 2 and Fig. 3).

As mentioned above, due to the greater weight of the animals at measurement 2, more isoflurane was needed to maintain a comparable level of anesthesia and respiration rate (Supplementary Table 1). Since the vasodilatory action of high isoflurane doses has an unfavorable effect on the magnitude of BOLD responses ^22^, direct comparison or pooling of the measurements would be inappropriate. Therefore, we regard measurement 2 merely as a confirmatory measure.

### Decreased cerebellar Purkinje cell number revealed by calbindin D28k immunostaining

Because lower Purkinje cell density is one of the most reproduced cerebellar postmortem anatomical findings in ASD, we made calbindin D28k (CB) immunostained sagittal sections from the cerebellums of 27 rats (9 from each group). We found that the number of CB-positive Purkinje cells was significantly lower in both VPA groups compared to the VEH group (Fig. 4) in all lobules tested (6a, 6b, 6c, 7). The difference was a larger magnitude in the VPA400 group. All p-values were smaller than 0.005, implying very strong evidence of Purkinje cell loss. In all lobules, the magnitude of the Purkinje cell number decrease was stronger in the VPA400 group (−30.7%, −47.9% −29.2% and −32.6% in lobules 6a, 6b, 6c and 7, respectively) than in the VPA600 group (−21.4%, −19.5%, −26.3% and −24.3% in lobules 6a, 6b, 6c and 7, respectively). The staining intensity remained normal. Summary statistics of normalized Purkinje cell numbers found by calbindin D28k immunostaining are listed in Supplementary Table 6.

**Figure 4.**
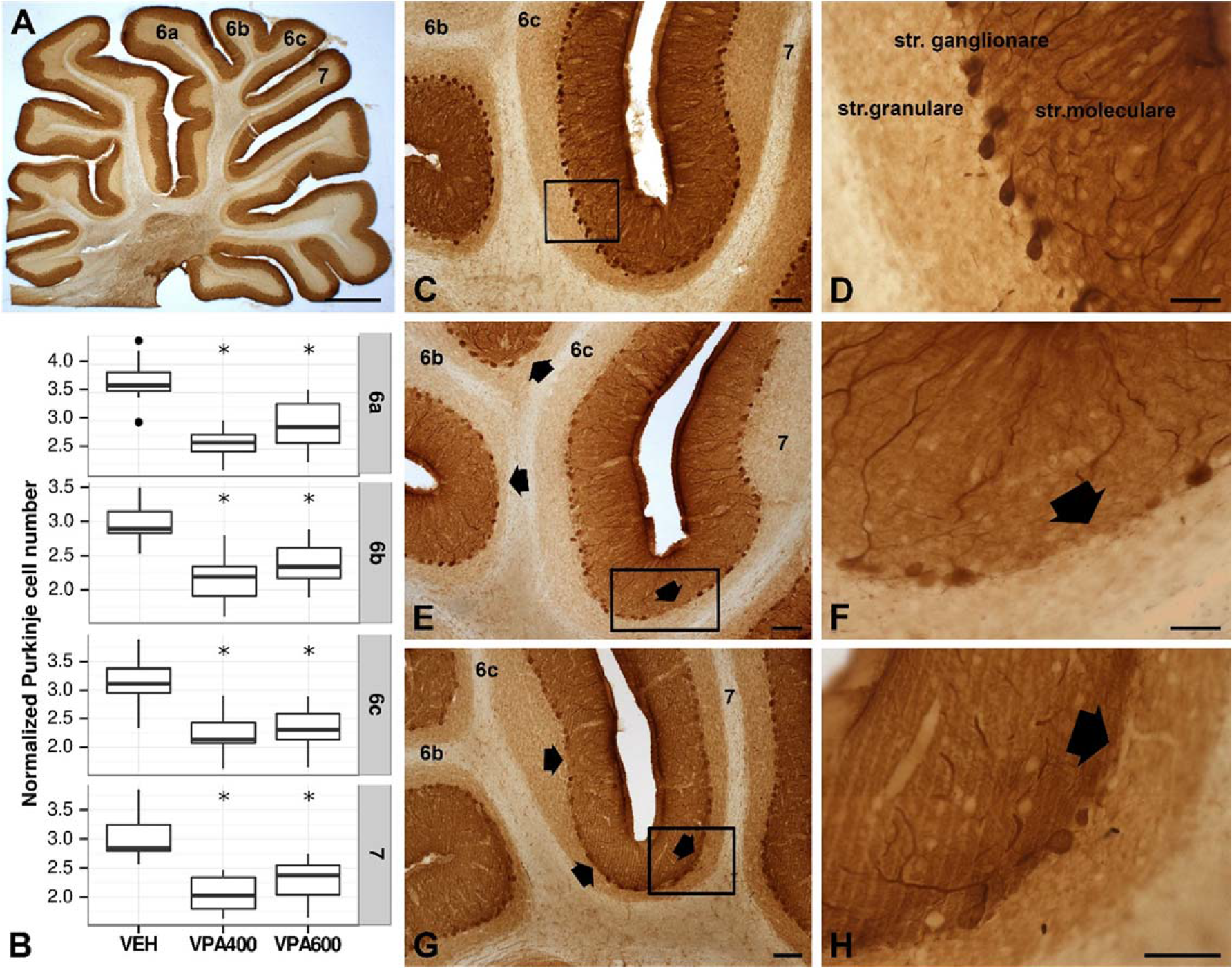
Prenatal valproate exposure induced reductions in cerebellar Purkinje cell number. (A) Light micrograph of calbindin D28k (CB) immunostained sagittal section of rat cerebellum at low magnification. (B) Boxplots representing normalized CB immunoreactive Purkinje cell numbers in cerebellar lobules 6a, 6b, 6c and 7, for the VEH, VPA400 and VPA600 groups. Asterisks depict significant (p<0.05) decreases compared to the VEH group. (C, D) Normal distribution of CB immunoreactive Purkinje cells in the ganglionic layer of cerebellar cortex on VEH sections. (E,F,G,H) Decreased CB-positive Purkinje cell number, found both in the VPA400 and VPA600 groups, while the staining intensity remained normal. The arrows point to unlikely large areas with lack of Purkinje cells. Scale bars: 1000 um (A), 100 um (C, E, G), 50 um (H), 20 um (D, E).

### Correlation of the BOLD response with the cerebellar Purkinje cell number

To the best of our knowledge, it is not known how lower Purkinje cell number is related to abnormal brain function in the VPA-model of ASD. Here, we analyzed how the correlation of individual Purkinje cell number and BOLD response to somatosensory stimulation is related to prenatal VPA exposure. In the VPA400 group (in animals with histological data available), a significant positive correlation of BOLD response with cerebellar Purkinje cell number was found in the frontal and cerebellar areas (Fig. 5B, Supplementary Table 10). In contrast, the VPA600 group exhibited a significant negative correlation in cortical and cerebellar regions (Fig. 5C, Supplementary Table 11). No significant correlation was found in the control group (Fig. 5A).

**Figure 5.**
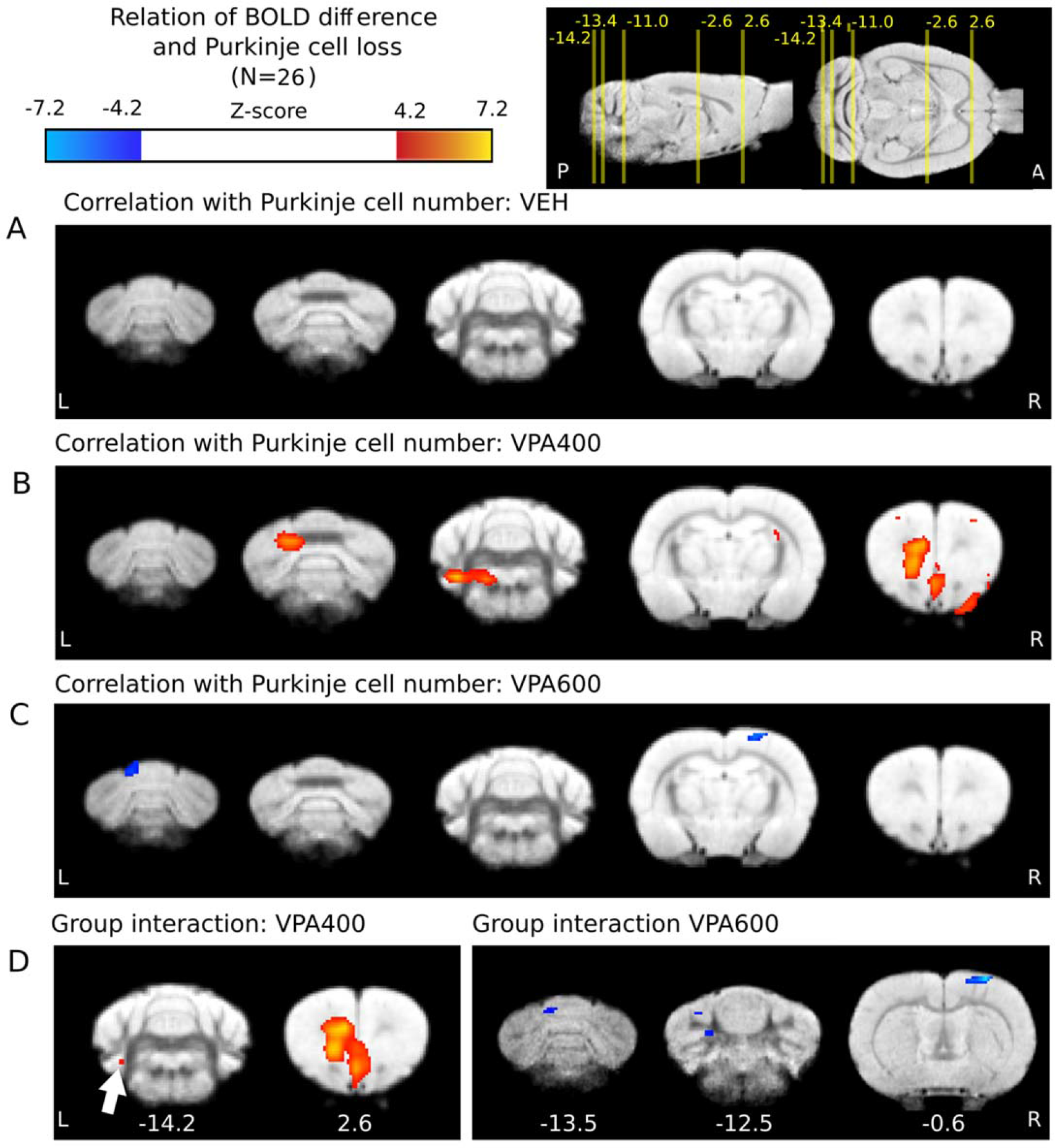
Prenatal valproate changes in the correlation of BOLD response and Purkinje cell number are present in both the cerebellum and the cerebral cortex. (A) No significant correlation of BOLD response with Purkinje cell number in the control group. (B) Significant positive interrelationship in the frontal and cerebellar areas in the VPA400 group. (C) Negatively correlated cortical and cerebellar regions with Purkinje cell number in the VPA600 group (D) Significant interaction effect between Purkinje cell number and treatment group on the BOLD response in a frontal and a cerebellar region of the VPA400 group (left side) and in somatosensory and cerebellar regions of the VPA600 group. Compared to controls, red indicates a greater regression slope in the VPA400 group, and blue denotes a lower slope in the VPA600 group. The distance of slices from bregma are shown under the slices. For effect sizes, see Supplementary Tables 12-13. Activations are displayed on coronal slices, overlaid on an in-house standard proton-density template ^23^. On the top right, the positions of the slices are shown on sagittal and horizontal planes as a distance from bregma in mm. Colorbars represent Z-score values. Significance thresholding was corrected for multiple comparisons (p=0.05). Abbreviations: A: anterior, P: posterior, L: left, R: right.

### Interaction effect of cerebellar Purkinje cell number and valproate treatment on the cerebellar and cerebral BOLD response

To capture VPA exposure-related effects, the differences in the groupwise “BOLD-Purkinje” correlation maps were further investigated by an interaction analysis. This revealed that the regression slopes between BOLD response and Purkinje cell number in the medial frontal cortex and in a small cerebellar region are significantly larger in the VPA400 group than in the vehicle group (Fig. 5D). We also observed areas of significant interaction in the VPA600 group. Here, the interaction coefficient was negative, implying that the regression slopes between BOLD response and Purkinje cell number in some cerebellar and somatosensory regions are significantly smaller (in fact, more negative) in the VPA600 group than in the vehicle group (Fig. 5D). The significant cerebral interaction between VPA treatment and Purkinje cell number suggests a VPA-induced cerebrocerebellar malfunction that is unseen when only focused on the cerebellum.

To gain a more direct insight into the *treatment × Purkinje* interaction effect on BOLD response, we conducted an ROI analysis. We defined two unbiased ROIs and investigated the effect of cerebellar Purkinje cell number on the individual ROI-mean cerebellar and cerebral BOLD responses. Analysis of the cerebellar ROI (left part of Fig. 6) revealed a general positive correlation between cerebellar BOLD response and Purkinje cell number in the VEH group. No such correlation was observed in the VPA600 group; however, the values might represent the naturally continuing trajectory of the relationship seen in the VEH group, but in ranges of the lower Purkinje numbers, where the BOLD response already converges to zero. In contrast, the strong positive correlation in the VPA400 display no continuity to the VEH group. In this case, despite the remarkable decrease in Purkinje cell number, the BOLD responses were comparable to those of the controls.

**Figure 6.**
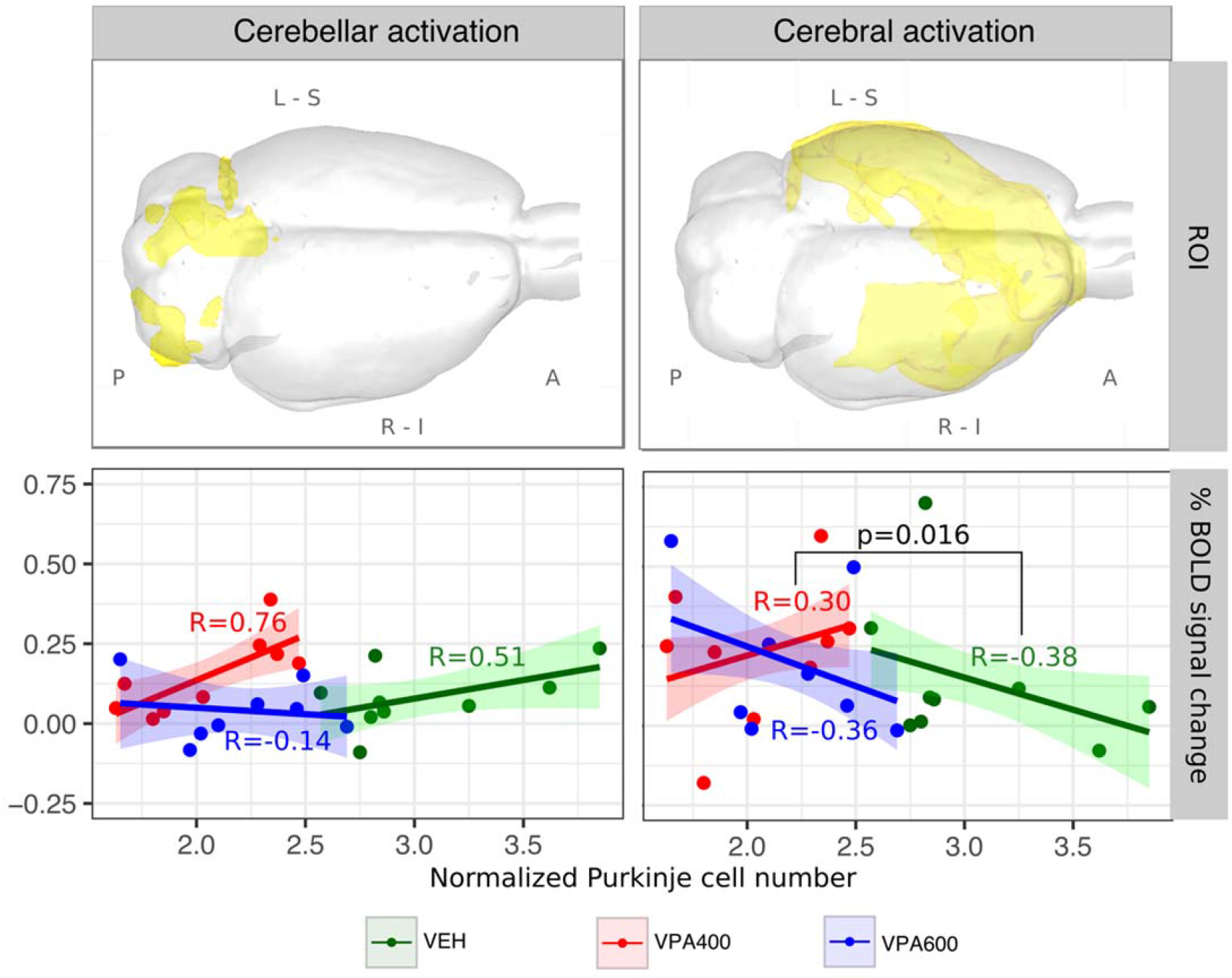
Atypical relationship between BOLD response and Purkinje cell number in cerebellar and cerebral ROIs of the pooled activation maps, suggesting probable malfunction of reciprocal cerebrocerebellar circuits. ROIs (visualized in 3D glass-brains) were defined based on the F-contrast representing the unbiased, pooled activation map, thresholded at FWER p<0.05 and divided into cerebellar (left) and cerebral (right) components. Analysis of the cerebellar ROI (left column) revealed a general positive correlation between frontal BOLD response and Purkinje cell number in the VEH group. No such correlation was observed in the VPA600 group. Despite the remarkable decrease in Purkinje cell number in the VPA400 group, BOLD responses, in terms of both magnitude and correlation with Purkinje cell number, were comparable to those of the controls. In the cerebral ROI (right panel), both the VEH and the VPA600 groups exhibited a significant negative correlation between BOLD response and Purkinje cell number. Interestingly, the correlation was positive in the VPA400 group, which possibly indicates the presence of an atypical cerebrocerebellar circuit in this VPA group. Abbreviations: A: anterior, P: posterior, L: left, R: right, S: superior, I: inferior.

In the cerebral activation areas (right side), both the VEH and the VPA600 groups exhibited a significant negative correlation between BOLD response and Purkinje cell number. However, this relationship became positive in the VPA400 group. The difference in slopes was captured by a significant interaction effect. Despite the lower BOLD responses due to anesthesia-related issues with larger animals, the second MRI measurement confirmed the tendency of a positive correlation between BOLD response and Purkinje cell number (Supplementary Fig. 5)

## Discussion

Although autism has a strong genetic component, there are many cases of autism, termed “idiopathic”, that are likely influenced by environmental factors ^24^. Animal models of idiopathic ASD include either inbred rodent strains that mimic ASD behaviors or models developed by environmental interventions such as prenatal exposure to VPA ^25^. This model of ASD exhibits face, construct and predictive validity and properly represents the epigenetic origin of idiopathic ASD ^26, 27^. While a few reports have already examined cerebellar anomalies in the prenatal VPA rat model ^28, 29^, the relationship of the abnormal Purkinje cell number to the function of cerebrocerebellar circuitry has not yet been investigated.

### ASD-like cerebral and cerebellar structural changes

In our study, brain hypoplasia was globally detected in the VPA600 group. In preclinical studies, even higher prenatal dose regimens are often used with similar results ^29, 30^; however, the observed magnitude of whole brain volume differences might be somewhat severe. On the other hand, the smaller dose (VPA400 group) by still inducing a focal, cerebellar gray matter difference (Fig. 1) might be in an appropriate range for triggering autistic-like morphological features without causing severe hypoplasia. Moreover, cerebellum-specific morphometric alterations, which could be a signature of human ASD, were only detected with this prenatal dose. These morphometric alterations localized in the cerebellar crus I/II and paraflocculus (in both measurements) compared to the VEH control group and the VPA600 group. The paraflocculus could be involved in atypical gaze ^31^, the posterior cerebellar vermis (vermal lobules VI-VII) and right crus I in repetitive behaviors and stereotyped interests in ASD ^32^. Crus I/II abnormalities are related to more severe ASD impairments in all domains ^7^ with altered functional connectivity to specific cerebral regions^11, 33, 34^.

Our structural MRI findings in this model may contradict the widely known “early brain overgrowth” phenotyping efforts in ASD ^35–37^, although increases in brain volume in high-risk infants may co-occur with excessive extra-axial cerebrospinal fluid (CSF)^5^. Nonetheless, several studies have suggested divergent age effects on brain morphology changes in ASD ^36^, with a more rapid age-related cortical thinning in temporal and parietal regions compared to controls ^38^. Decreased thickness in the temporal cortex was reported recently from a large multinational sample of the ENIGMA ASD working group, while at the same time, increased cortical thickness in the frontal cortex was found ^39^. Similarly, using data from the multicenter repository, the ABIDE initiative, a surface-based cortical thickness analysis revealed an increase in prefrontal cortical thickness in ASD ^40^. Remarkably, an inverse correlation had been found earlier between the sizes of the frontal lobe and cerebellum in ASD ^41^, reflecting an abnormal overgrowth in the frontal lobes ^42^.

### Sensory BOLD hyperactivation induced by prenatal valproate exposure

Sensory abnormalities are known in ASD (DSM-5, diagnostic criteria B.4.) Somatosensory evoked potentials show hyperactivity in infantile autism ^43^ as in Fmr1(−/y) mice ^44^; therefore, sensory hypersensitivity may have translational value. Prenatal VPA rats have also shown increased sensory reactivity to mechanical stimuli ^45^ and increased neuronal activation in the auditory brainstem after sound exposure ^46^. Interestingly, local functional overconnectivity was not demonstrated in the somatosensory cortex of ASD individuals by magnetoencephalography, challenging the theory that the primary substrate of hypersensitivity is localized in the somatosensory cortex ^47^.

In our fMRI measurement, the presence of altered sensory processing was reinforced in the prenatal VPA model of autism. The BOLD hyperactivation was more pronounced in the VPA400 group, but it was also clearly present in the VPA600 group. BOLD hyperactivation was prominent in areas with a significant group-mean response in the VEH group, and frontal hyperactivation, including the lateral and ventral orbital, prelimbic, frontal association, and anterior cingular cortices, was also found in the VPA groups. In the VPA400 group, there were additional activations in the brainstem and cerebellar regions (specifically in the crus II); however, these activations proved to be significant only when compared to the VPA600 group.

Although both the hypoplasia found in our VBM analysis and the cerebellar hyperactivations in the VPA400 group were localized in nonsensorimotor areas of the cerebellum (paraflocculus and crus I/II), a direct link of structural changes to the observed sensory hyperactivation might also be possible, since there is evidence for a noncanonical organization of cerebrocerebellar circuits in ASD, especially in the sensory domain ^48, 49^. Since the paraflocculus has projections to the brainstem and the crus I/II normally interconnects with frontal and parietal areas, the brainstem (lemniscus) and frontal components of the BOLD hyperactivation might also reflect the consequences of noncanonically organized cerebrocerebellar circuits. Consistent with this interpretation, abnormal cortical connectivity of the crus I/II region has been found in ASD ^7, 8, 11^.

### Atypical effect of decreased Purkinje cell number on cerebrocerebellar activity

Following current ideas about the atypical functional connectivity of cerebrocerebellar circuit loops in ASD ^7, 8, 48, 49^, it is straightforward to hypothesize that the significant VPA-induced decrease in Purkinje cell number might have a developmental effect on the cerebral cortical regions the cerebellum functionally connected with ^8, 20^. We addressed this assumption by testing the groupwise correlation of BOLD response and Purkinje cell number and whether it was different between groups (evaluating the “*treatment × Purkinje cell number”* interaction effect on the BOLD response).

In the case of the cerebellar BOLD signal, the main contributor to the activity is the parallel / mossy fiber system ^50^, and the contribution of the Purkinje cell activity might be relatively small ^51^. Therefore, instead of simply reflecting the summed activity of Purkinje cells, the positive correlation between BOLD response and Purkinje cell number observed in the control group might be driven by the indirect effect of increased cerebral input through the parallel / mossy fiber systems.

On the other hand, throughout the cerebral activation areas, the BOLD - Purkinje cell number correlation is negative in the control group, which might be either a consequence of a stronger modulatory / inhibitory effect in the presence of more Purkinje cells or might be caused by some neurodevelopmental source affecting both Purkinje cell development and the maturation of distant cerebral circuits. Indeed, on the Purkinje cell level, prenatal VPA exposure delayed the developmental excitatory / inhibitory GABA “switch” ^52^, supporting a role of developmental timing in the pathophysiology of ASD.

The correlation of the BOLD response and Purkinje cell number is strongly dependent on the dose of VPA exposure. The simplest explanation for the significant “*group × Purkinje cell number”* interaction in the VPA600 group is that they represent a naturally continuing, nonlinear trajectory of the VEH group but in ranges of significantly lower Purkinje cell numbers, where the BOLD response already saturates to zero (thereby introducing a difference in linear regression slopes; also visible in Fig. 6). However, this explanation does not hold for the VPA400 group. Here, our results suggest that, despite the lower Purkinje cell numbers, the BOLD-Purkinje correlation maintained at similar or even higher levels than those of the control group, possibly by means of some compensatory mechanisms. Importantly, this “compensated” cerebellar BOLD response is accompanied by an atypical effect on cerebral responses, where the BOLD-Purkinje correlation turns from negative to positive. Notably, the atypical BOLD-Purkinje correlation is most pronounced in the frontal activation areas, supporting our hypothesis that noncanonically organized cerebrocerebellar circuits are present between the crus I/II and frontal regions. This might be a sign of a drastic VPA-related change in cerebrocerebellar modulation and/or a consequence of VPA-induced developmental diaschisis ^20^. While the underlying mechanism remains unclear, our results demonstrate that measurable correlations exist in the VPA model between the cerebellar morphology and cerebrocerebellar circuit loop function.

## Conclusion

In this multimodal MRI study of the VPA rat model of ASD, we found (i) ASD-like global and cerebellar changes in brain morphology, (ii) a functional BOLD hyperactivation to whisker sensory stimuli and (iii) in the combined analysis of altered cerebellar histology and BOLD responses, the presence of atypical cerebrocerebellar circuits. Our results suggest an altered interaction between the cerebellar and frontal cerebral areas. These alterations in the rat VPA model of ASD have potential translational value in the search for objective neuroimaging biomarkers in ASD.

## Methods

### Subjects

Timed-mated female Sprague-Dawley rats (Harlan, San Pietro al Natisone, Italy) arrived at the local vivarium at gestational day 6-8. Animals were kept in polycarbonate cages in a thermostatically controlled room at 21±1°C. The room was artificially illuminated from 6 a.m. to 6 p.m. The rats were fed conventional laboratory rat food (sniff R/M + H Spezieldiäten GmbH D-59494 Soest). All of the procedures conformed to the guidelines of the National Institutes of Health for the care and use of laboratory animals, were approved by the Local Ethical Committee of Gedeon Richter Plc. and were carried out in strict compliance with the European Directive 2010/63/EU regarding the care and use of laboratory animals for experimental procedures. All efforts were made to minimize the number of animals as well as their suffering. Experiments were performed and reported according to the ARRIVE guidelines on animal research.

### Gestational valproic acid administration

The timed-pregnant female Sprague-Dawley rats received a single subcutaneous dose of 400 or 600 mg/kg sodium VPA (Sigma-Aldrich Chemical Co, UK) or vehicle (saline injection) in the loose skin area of the neck in a volume of 1 ml/kg dissolved in sterile physiological saline at gestational day 12.5 as described previously^53^. The treatment groups are referred to as VEH, VPA400 and VPA600. On postnatal day 21, male offspring were weaned and housed in groups of 3 or 4.

### Image acquisition

Before magnetic resonance imaging, animals were habituated to the acoustic noise of the MRI scanner for 1 week in a separate animal room. In the experiments, 36 male rats (12 rats from each treatment group) were used. The mean (± standard deviation) initial weight of the rats was 209 (±27) g over the first MRI measurements (performed in a period of two weeks) and 393 (±23) g over the second MRI measurements (beginning one month after the end of the first measurement). Four rats were scanned per day in a random order from each treatment group.

On the day of scanning, rats were anesthetized before transporting to the magnet room. Anesthesia was introduced with 5% isoflurane and then maintained at approximately 1.25% during scanning. The body temperature was maintained at 38±1°C with thermostatically controlled air flow around the rat. The respiration of the animal was monitored continuously with a small pneumatic pillow sensor during the experiment (SA Instruments, Inc., NY, USA). The MRI experiments were performed using a 9.4 T Varian MRI system (Varian Associates Inc., Palo Alto, CA) with a free bore size of 210 mm, which contained a 120 mm inner size gradient coil (minimum rise time 140 μs; 200 μs were used). For excitation, an actively RF-decoupled 2 channel volume coil system with an inner size of 70 mm was used, and a fixed-tuned receive-only phase array rat brain coil (RAPID Biomedical GmbH, Rimpar, Germany) was located directly above the dorsal surface of each rat’s head to maximize the signal-to-noise ratio.

For the spatial coregistration of fMRI scans and for voxel-based morphometry (VBM), proton density weighted anatomical scans were acquired using gradient echo multi-slice imaging (GEMS; echo time (TE): 5.78 ms, repetition time (TR): 855 ms, flip angle: 40^◦^, data matrix: 256 × 256, total scan time: 22 min 30 s) sequence with a field-of-view (FOV) of 40 mm × 40 mm and a slice thickness of 0.2 mm, without interslice gap. Seventy-four slices were acquired in an interleaved order. Anatomical scans were repeated 8 times and then averaged into a single image.

### MRI measurements: anesthesia, respiration and motion

Since motion artifacts are very problematic in MRI, before any further analysis, we examined the magnitude and between-group differences in in-scanner motion, respiration and anesthetic dose. Despite the superficial anesthesia during functional MR imaging, motion was moderate throughout all MRI experiments (Supplementary Table 1). For both MRI measurements, weight, average in-scanner motion (as estimated by the motion correction algorithm), isoflurane dose and respiration rate were not different between the groups. Summary statistics of these measures are listed in Supplementary Table 1.

### Functional MRI experiments

The T2*-weighted echo planar imaging (EPI) sequence for the fMRI experiments had the following settings: TE: 10 ms, TR: 2030 ms, flip angle: 90^◦^, averages: 1, dummy scans: 4, data matrix: 50 × 50, 400 repetitions, FOV: 40 mm × 40 mm, slice thickness: 0.8 mm, no interslice gap, and 16 horizontal slices (resolution: 0.8 × 0.8 × 0.8 mm^3^). For the reduction of the EPI Nyquist ghost artifact, a triple reference scan^54^ was used, meaning that the functional images were acquired with two opposite gradient polarities, which resulted in an effective TR of 4060 ms.

For a somatosensory stimulus, pneumatic “air-puffed” whisker stimulation was used in a block-design fashion. The stimulation was delivered through a tubing system that was integrated into the holding cradle. The air pressure was set to 1 bar. Air puffs were delivered at a frequency of 1 Hz with a blowing time of 200 ms. The duration of the stimulation blocks was 30 s followed by a 60 s rest. Over one experimental session, 6 blocks of stimulation were used. The pneumatic stimulation was controlled by a custom-programmed user interface developed in LabView (National Instruments Corporation, Austin, TX, USA).

### Exclusion criteria

Exclusion criteria were fixed prior to data analysis as follows: scans with striking imaging artifacts (based on visual inspection) or with an average root mean squared relative displacement greater than 0.05 mm during the fMRI scan (as calculated by FSL) were subject to exclusion from all analyses.

Moreover, subjects without immunohistological data were excluded from the analyses focusing on the correlation of BOLD response with Purkinje cell number and its group-level interaction.

None of the scans had to be discarded due to artifacts or in-scanner motion issues, resulting in N=36 for the whole-brain fMRI analysis of the first measurement. On the other hand, in the second measurement, 3 animals had to be excluded because of in-scanner motion and inadequate anesthesia (2 controls and 1 in the VPA600 group), resulting in N=33.

The immunohistology experiments were only performed for N=9 animals per group, and one animal from the VPA600 group had to be excluded for technical reasons.

### Imaging data analysis

To obtain maximally translatable results, we followed the conventions set for human MRI analysis, similar to our previous works^23, 55^. Small-animal imaging-specific features of the applied image processing pipeline are described in detail. When not specified otherwise, the same data analysis steps were performed in the first and second MRI measurements.

### Preprocessing

The raw images were converted to NIfTI-format (Neuroimaging Informatics Technology Initiative) by an in-house developed script. The anatomical and functional images were reoriented to match the standard orientation of the digitalized version of the Rat Brain Atlas of Paxinos and Watson^56^. All images were rescaled by a factor of ten to achieve image dimensions similar to human data and thus facilitate the use of image processing algorithms developed for human image analysis. All the approaches performed on the upscaled images were scale-invariant, except for a built-in constraint optimization for motion correction. The upscaling only affects the dimension descriptor fields in the file header and does not involve interpolation or any information loss. The analysis was carried out in a multistage process using the image analysis software package, FSL^57^ (FMRIB’s Software Library, www.fmrib.ox.ac.uk/fsl), and in-house developed software tools. All the fMRI time-series were motion-corrected using MCFLIRT^58^. FSL BET (Brain Extraction Tool^59^) was used to remove nonbrain areas from the structural and functional images. The fractional intensity threshold was set to 0.65 and 0.7 for functional and structural images, respectively, and the vertical gradient in the fractional threshold was set to 0.1. Before the brain extraction, the images were rescaled in the y-direction by a factor of 0.5 to ensure that the spherical brain model used by BET gained robust segmentation results. The brain-extracted images were then rescaled to the original y-axis dimensions.

To achieve spatial correspondence for the group analysis, all images were spatially standardized using FSL FLIRT^60^, which utilized a 6-parameter rigid-body transformation. The high-resolution structural image was fitted to an in-house developed standard template^23^ (average of 200 nonlinearly fitted structural images) by an affine transformation (FLIRT) and a nonlinear deformation field estimated by FNIRT^61^. In the latter coregistration procedure, a 10 × 10 × 10 mm^3^ warping field (in upscaled space) was estimated with three iterations using a relatively conservative lambda parameter set of 60, 40 and 20. This stricter regularization was applied as the morphological diversity of the rat brain is smaller compared to humans.

### Voxelwise functional MRI (fMRI) analysis

Before the statistical analysis, the functional images were spatially smoothed using a Gaussian kernel of 12.0 mm FWHM (note that the images were upscaled). High-pass temporal filtering (Gaussian-weighted least-squares straight line fitting, with sigma=140.0 s) was then applied to remove slow drifts from the signal. The fMRI data processing was carried out using FEAT (FMRI Expert Analysis Tool) Version 6.00, part of FSL. After signal prewhitening (FILM), the first-level statistical analysis on the time-series was carried out using the general linear model with local autocorrelation correction^62^. The analysis modeled each individual rat’s data from each session and included 7 explanatory variables in total: one regressor modeling the stimuli (convolved with a double-gamma canonical HRF), its temporal derivate, to account for slight differences in timing and 5 noise-ROI-based, CompCor^63^ confounder variables. For each fMRI session, a noise ROI was delineated based on the temporal signal-to-noise ratio of the BOLD signals (the upper 2 percentiles of within-brain voxels were chosen on each slice), and the first five principal components of the corresponding time-series were extracted from the data, following the broadly used t-CompCor technique^63^. Individual statistical Z-score images for the first regressor were obtained (average BOLD response to the stimuli).

Using the spatial transformations computed during image coregistration and standardization, Z-score maps resulting from the individual statistical analysis were realigned to the common standard space to achieve spatial correspondence during the group-level voxelwise analysis.

A second-level general linear model analysis was performed with FEAT. Data from the first and second measurements were analyzed separately. Three different statistical models were evaluated for both measurements.

***Model 1*** was a simple model with three explanatory variables modeling the treatment groups with dummy coding (N=36 and 33 for measurement 1 and 2, respectively). Statistical contrasts were defined to assess the VPA400 vs. VEH., VPA600 vs. VEH and VPA400 vs. VPA600 comparisons. An F-contrast was also specified to establish a statistical map of voxels activated in at least one of the three groups. This map was used for defining unbiased ROIs.

***Model 2*** was a model analogous to Model 1 but with the restricted population for which histological data were also available (N=25, for both measurements). The goal of this model was simply to demonstrate that the results of model 1 are not driven by animals excluded from further analysis (that is, the restricted population is still representative in terms of findings in the whole population).

***Model 3***, next to the three explanatory variables of Model 2 (representing the VPA groups), involved three extra explanatory variables. Those represent the statistical interaction, by containing the demeaned (globally, for all included subjects), normalized Purkinje cell numbers of the cerebellar lobule 7, multiplied by the group-level explanatory variables, respectively. With the proper contrasts, this model allowed for analyzing the groupwise correlations between Purkinje cell number and BOLD response, as well as the interaction effects, comparing the regression slopes between groups (N=25 for both measurements).

The resulting group-level statistical maps were corrected for multiple comparisons on the voxelwise-level, with a GRF-theory-based maximum height thresholding procedure and thresholded with a significance threshold of p=0.05 ^64^. For the second measurement, a cluster-based thresholding was also applied, with a Z=3.1 cluster-forming threshold and a cluster probability cutoff of p=0.05. This thresholding method, although less effective in localizing the activation, was used because the voxelwise thresholding was too conservative for the observed lower effect sizes in the group-contrast images of measurement 2.

### ROI analysis

In addition to voxel-based analyses, a cerebral and cerebellar ROI was defined, and the individual ROI-mean percent BOLD signal change was calculated and compared between groups. These ROIs were delineated based on the unbiased F-contrast of Model 1 to avoid favoring any of the groups. This activation map was thresholded with a GRF theory-based maximum height thresholding procedure and thresholded with a significance threshold of p=0.05 to identify voxels that were activated in at least one of the groups. The resulting activation mask was than multiplied by in-house cerebellum and cerebrum masks to define the cerebellar and cerebral ROIs of activation, respectively. These were used for the ROI-level interaction analysis (Fig. 6).

### MRI volumetric and voxel-based morphometry analysis

We used the voxel-based morphometry (VBM)-style protocol^65^ of FSL^59^, but optimized for the rat brain, as in our previous study ^55^. Nonbrain parts were removed from all the structural images ^59^, and tissue-type segmentation was carried out by FAST4^66^. To obtain optimal tissue classification, an in-house *a priori* tissue map ^55^ was used. The volume of whole-brain gray and white matter was estimated by taking the sum of the corresponding partial-volume estimate maps.

The resulting gray-matter partial volume images were registered to a standard space using linear transformation^58^, followed by a nonlinear registration^61^. The resulting images were averaged to create a study-specific template, to which the native-space gray matter images were then nonlinearly reregistered. The registered partial volume images were then modulated (to correct for local expansion or contraction) by being divided with the Jacobian of the warp field (incorporating the linear scaling factor, as well). The modulated segmented images were then smoothed with an isotropic Gaussian kernel with a sigma of 4 mm. Finally, statistical inference on the treatment effect was performed by voxelwise GLM with permutation-based nonparametric testing (FSL’s randomise). Cluster-based belief-boosting was carried out by the TFCE technique^67^. The images were then thresholded with permutation-based empirical maximum height thresholding at a (corrected) significance threshold of p=0.05^64^.

### Histological processing

#### Perfusion and preparation of tissue sections

The animals were deeply anesthetized using intraperitoneal injection of an anesthetic mixture (containing ketamine, xylazine hydrochloride, promethazinium chloride) and perfused through the heart first with saline (2–3 min) followed by a fixative (30 min) containing 4% paraformaldehyde in 0.1 M phosphate buffer (PB) (pH 7.4). Brains were removed from the skull. After vibratome slicing and extensive washing in 0.1 M PB, the 60 μm-thick sections were incubated in 30% sucrose overnight, followed by freeze thawing over liquid nitrogen four times.

#### Immunocytochemistry

The sections were processed for immunoperoxidase staining. All washing steps and dilutions of the antibodies were performed in 0.05 M Tris-buffered saline (TBS), pH 7.4. After extensive washing in TBS, the sections were blocked in 3% bovine serum albumin for 45 min and then incubated in rabbit anti-calbindin-D28kD antibody (1:1000, Swant) for a minimum of 48 h at 4°C. Following the primary antisera, the sections were treated with biotinylated anti-rabbit IgG (1:300) raised in goat for 2 h and then with avidin biotinylated horseradish peroxidase complex (1:500; Elite ABC; Vector Laboratories) for 3 h at room temperature. The immunoperoxidase reaction was developed using 3,3’-diaminobenzidine (DAB) as the chromogen.

### Statistical analysis of brain volumetry, ROI and histological data

Gray and white matter volumetry, cerebral and cerebellar ROI and lobule-wise histological data were compared by means of permutation testing to overcome possible normality issues. Permutation-based linear models (R-package lmPerm) were fitted to explain these data with a factor variable representing the treatment groups. Additionally, for the interaction analysis of the cerebral and cerebellar ROIs, the demeaned, normalized Purkinje cell number of the cerebellar lobe 7, and its interaction with the group factor variable was also added.

The number of maximum permutations was set to 100000, and factor variables were explicitly set to be handled with the “treatment” contrast of R (traditional “dummy coding”). Permutations were stopped when the estimated standard error of the estimated p was less than 0.01*p.

### Data availability statement

Data generated and analysed during this study are included in this published article and its Supplementary Information material, the full datasets generated during and/or analysed during the current study are available from the corresponding author on reasonable request.

## Acknowledgements

The authors are thankful to Prof. Dagmar Timmann-Braun for her most valuable insights regarding the interpretation of the cerebrocerebellar aspects of our results.

The authors are also grateful for the technical assistance of Anita Bérces, Tiborné Gyarmati, Pálma Diószegi and Katalin Tóthné Fekete. This work was supported by the Hungarian National Research, Development and Innovation Office (KMOP-1.1.5-08-2009-0001, ERNYO-13-1-2013-0003).

## Author Contributions Statement

TS, VR, GÉNy, GyL, LB, EP and ACz are designed the experiments. RK, KS, CKCs, AV, DG and EP are performed the experiments. TS, VR, GÉNy, ZsS, EP and ZsTK are analysed the data. TS, VR and ACz are wrote the main manuscript text and TS and EP prepared figures. All authors reviewed the manuscript

